# The Linear Element is a Stable Structure Along the Chromosome Axis in Fission Yeast

**DOI:** 10.1101/2020.07.03.185785

**Authors:** Da-Qiao Ding, Atsushi Matsuda, Kasumi Okamasa, Yasushi Hiraoka

## Abstract

Chromosomes structure changes dramatically upon entering meiosis to ensure the successful progression of meiosis-specific events. During this process, a multilayer proteinaceous structure called synaptonemal complex (SC) is formed in many eukaryotes. Instead, in the fission yeast *Schizosaccharomyces pombe*, linear elements (LinEs), which are structures related to an axial element of SC, form on the meiotic cohesin-based chromosome axis and are required for the formation of DNA double-strand breaks. In contrast to the well-organized SC structure, LinE structure had been observed only by silver-stained electron micrographs or in immuno-fluorescence stained spread nuclei. Thus, their fine structure and dynamics in intact living cells remain to be elucidated. In this study, we performed live cell imaging with wide-field fluorescence microscopy as well as 3D structured illumination microscopy (3D-SIM) for the four components of LinE, the Rec10, Rec25, Rec27 and Mug20. We found that LinEs consist of threads formed along the chromosome axes during the meiotic prophase. Rec10 binds to the chromosome itself and shapes into LinEs only in the presence of all the other LinE components. Rec25, Rec27, and Mug20 attach to the chromosome in the presence of Rec10. LinEs are stable in a short-time treatment with 1,6-hexanediol; and fluorescence recovery after photobleaching (FRAP) experiment reveals slow recovery from photobleaching, indicating a stable property of LinEs.

## Introduction

Sexual reproduction is accomplished through a special cell division called meiosis. Meiosis produces gamete cells through two rounds of consecutive chromosome segregation, which reduce the diploid set of homologous chromosomes in the gametes to a haploid set. One of the most important processes during meiosis is the recombination of homologous chromosomes coming from each of the parent cells. The successful recombination generates physical links between homologs and genetic variations in the offspring. Physical links called chiasmata are required for reductive segregation of homologous chromosomes during meiosis I.

In many organisms, a protein structure called synaptonemal complex (SC) forms in between and along the entire paired homologous chromosomes in meiotic prophase. The SC is composed of lateral elements that represent the axis of each homologous chromosome, a central element in between the lateral elements, and transversal elements that link the central element to the lateral elements. The SC facilitates the transformation of crossovers into chiasmata and the establishment of crossover interference in some organisms (Review in Zickler and Kleckner 2015).

Unlike many other organisms, the fission yeast *Schizosaccharomyces pombe* does not assemble canonical SC. Instead, it forms the so-called linear elements (LinEs) resembling evolutionarily related axial/lateral elements of the SC (Bahler et al. 1993; Lorenz et al. 2004; Loidl 2006). The LinE is composed of four essential components: Rec10, an ortholog of axial element protein Red1 in *Saccharomyces cerevisiae*, and three small coiled-coil proteins −Rec25, Rec27 and Mug20 (Lorenz et al. 2004; Davis et al. 2008; Spirek et al. 2009; Estreicher et al. 2012; Fowler et al. 2013). Rec10 and all the other LinEs components are required for DNA double strand break (DSB) formation and recombination (Lorenz et al. 2004; Davis et al. 2008; Estreicher et al. 2012; Fowler et al. 2013; Ma et al. 2017).

Meiotic cohesins are essential for sister chromatid cohesion. They are the main component of axial elements, which subsequently form lateral elements of SC (Page and Hawley 2003). Similar to SC, the establishment of LinEs requires a meiotic cohesin axis. In *S. pombe* meiosis, mitotic cohesins are replaced by meiotic cohesins Rec8 and Rec11 (Parisi et al. 1999; Watanabe and Nurse 1999; Yokobayashi et al. 2003). Also, a conserved cohesin-associated protein Pds5 is involved in the maintenance of the sister chromatid cohesion (van Heemst et al. 1999; Hartman et al. 2000; Panizza et al. 2000; Tanaka et al. 2001). In *S. pombe*, chromosomes become less compacted in the absence of Rec8 or Rec11, whereas the loss of Pds5 results in Rec8-dependent over-compaction (Ding et al. 2006; Sakuno and Watanabe 2015). Using three-dimensional super-resolution structured illumination microscopy (3D-SIM), we have shown that Rec8 forms a linear axis on chromosomes, which is required for the organized axial structure of chromatin during meiotic prophase; in the absence of Pds5, the Rec8 axis is shortened while the chromosomes are widened (Ding et al. 2015). LinE formation on chromosomes requires meiotic cohesin Rec8 and Rec11 (Molnar et al. 1995, 2003). Rec11 phosphorylation by casein kinase 1 recruits Rec10 to the chromosome (Sakuno and Watanabe 2015).

The fine structure of LinE has been examined by electron microscopy (EM), as well as immunofluorescence microscopy in spread cells. LinEs were observed as dots or short single lines at an early stage, then they become longer lines or bundles at late meiotic prophase (Bahler et al. 1993; Lorenz et al. 2004; Davis et al. 2008; Molnar et al. 2003). Live cell imaging of LinE components, tagged with GFP or other fluorescent proteins in intact cells, show dots like foci or linear structures (Fowler et al. 2013). Both EM and conventional live cell imaging remain problematic. First, the structure of LinE may not be well preserved or even reorganized during the nuclear spreading for EM specimen preparation. Second, the resolution of live cell fluorescence microscopy of is not high enough to reveal its fine structure. In this study, we examined the LinE using 3D-SIM aiming to elucidate its fine structure and dynamics in live meiotic cells. Our results show that LinEs are discontinuous threads formed along the chromosome axes during the meiotic prophase.

## Results

### LinE proteins cooperatively form the filamentous structure in living cells

To follow the dynamics of LinE during meiosis, we visualized LinE using GFP-tagged LinE components. The results are shown in Figure 1. Upon nitrogen starvation, haploid cells of the opposite mating types conjugate to form a zygote cell. Their nuclei fused with each other (karyogamy, “kar” in Fig. 1) to form an elongated diploid nucleus called the “horsetail” nucleus (Fig. 1). The horsetail nucleus represents the meiotic prophase in *S. pombe* (Chikashige et al. 1994; Ding et al. 2004). We found that all the LinE proteins appeared during karyogamy and disappeared at the end of the horsetail stage (Fig. 1). The GFP signals were not uniform but looked like many numerous uneven dashed lines. GFP signal intensity increased during the horsetail stage. All of the LinE proteins display undistinguishable staining pattern and dynamics (Fig. 1).

**Fig. 1.**
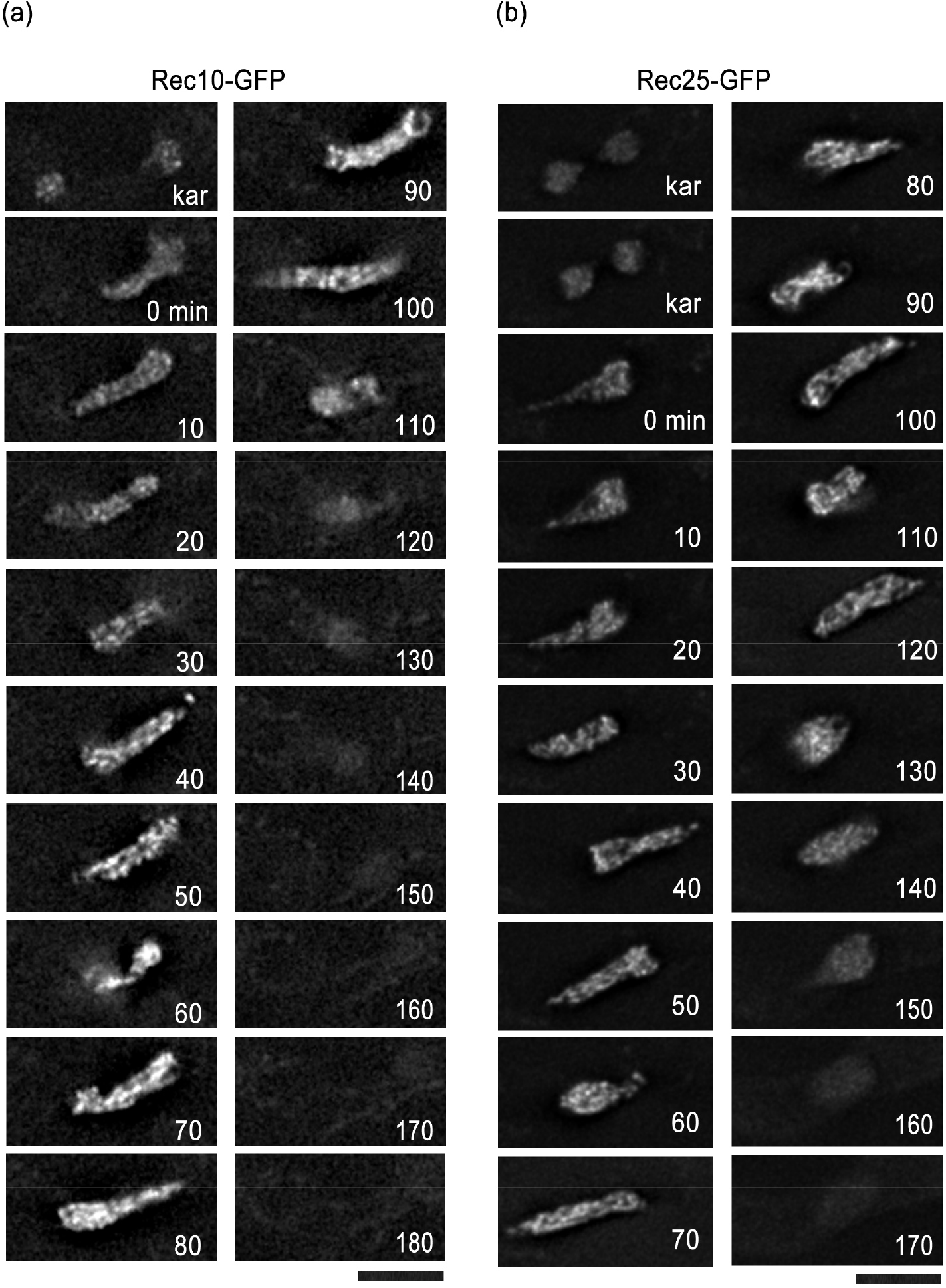
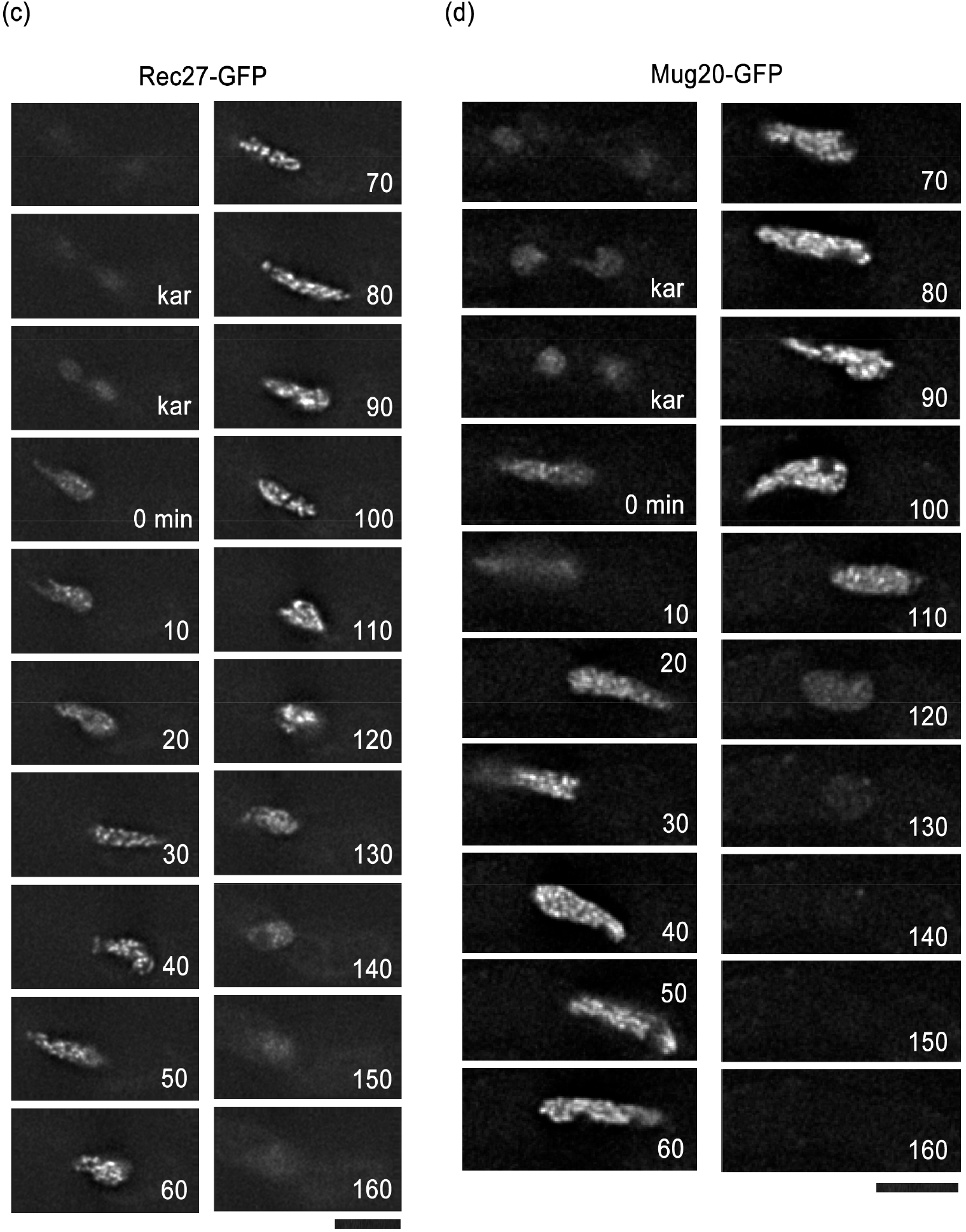
Dynamics of linE proteins in living cells during meiotic prophase. Selected time-lapse images of live cell observation of the LinE proteins in wild-type cells. (a) Rec10-GFP, (b)Rec25-GFP, (c) Rec27-GFP, and (d) Mug20-GFP. Labels “kar” stands for karyogamy. Numbers indicate the time in minutes after karyogamy. Each image is a single optical section from 3D deconvolved stacks collected at the indicated time point. Bar, 5 μm.

It has been shown that the loading of LinE components onto chromosomes are mutually interdependent (Davis et al. 2008; Fowler et al. 2013). To address their interdependency more clearly, we followed the formation of LinEs in various deletion backgrounds. In the absence of Rec25, Rec27 or Mug20, Rec10-GFP fluorescence was observed from the entire chromosomes, but they could not form LinEs-like linear fragments (Fig. 2a-c). The chromosomal localization of Rec10-GFP was also confirmed by 3D-SIM observation as shown below (see Fig. 5a). This Rec10 localization sustains to the first meiotic segregation (Fig. 2a, 2b, more than 190 min on average), which was never observed in wild-type cells (Fig. 1, about 148 min on average). This result suggests that Rec10 can form LinEs only in the presence of all the other LinE components. On the other hand, Mug20-GFP could not localized in the horsetail nucleus in a Rec10-deletion background (Fig. 2d), suggests that Rec10 is required for loading other LinE components onto the chromosome. In addition, in the absence of Rec25, Mug20-GFP signal was barely observed in the nucleus (Fig. 2e), which is consistent with a previous report (Fowler et al. 2013).

**Fig. 2.**
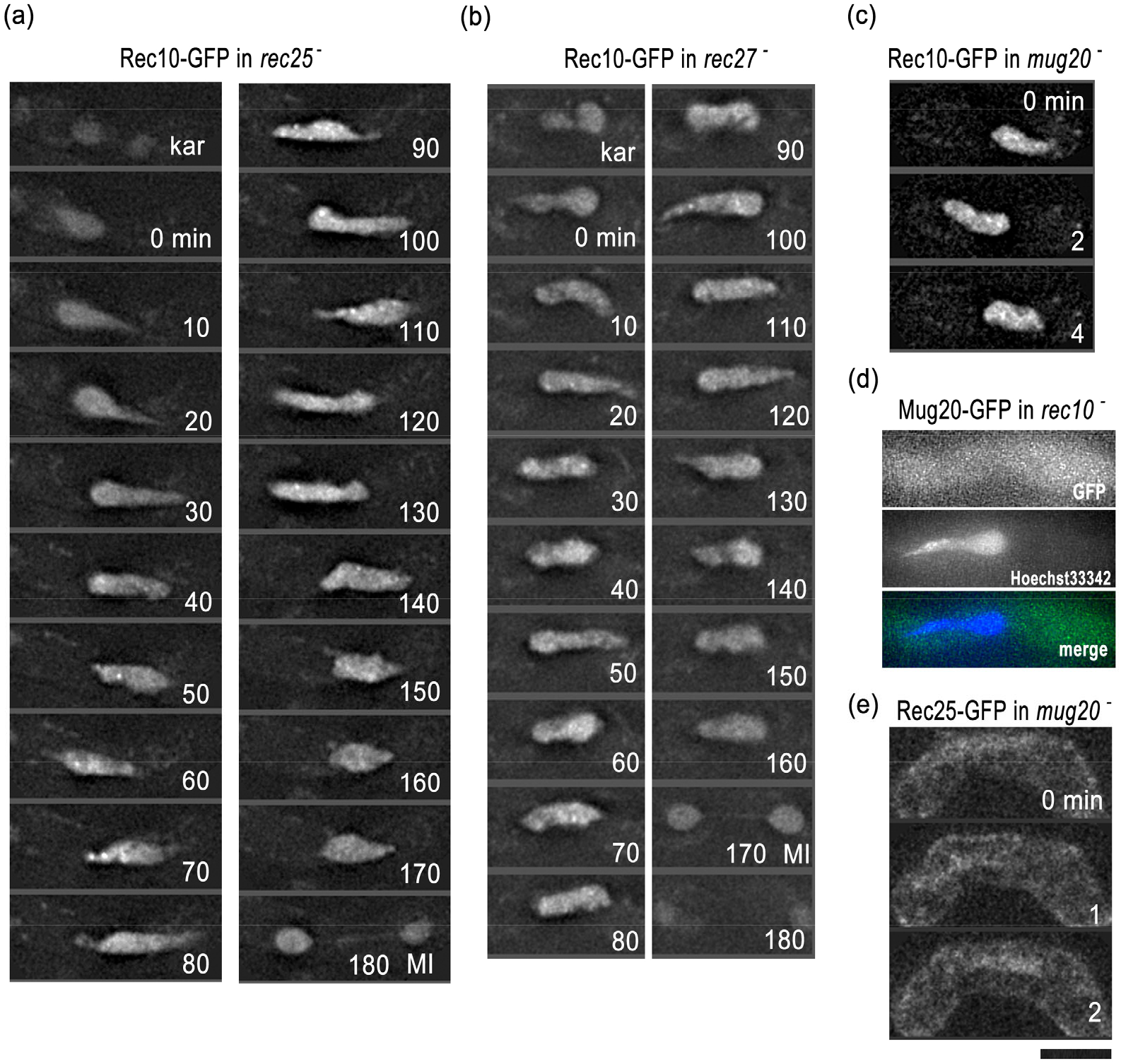
Interdependency of LinE components. Selected time-lapse images of live cell observations of the LinE proteins in various mutant backgrounds. (a) Rec10-GFP in a rec25-deletion cell. (B) Rec10-GFP in a rec27-deletion cell. (c) Rec10-GFP in a mug20-deletion cell. (d) Mug20-GFP in a rec10-deletion cell. The nucleus was stained with Hoechst33342. (e) Rec25-GFP in a mug20-deletion cell. Labels “kar” for karyogamy, “MI” for meiosis I. Each image is a single optical section from 3D deconvolved stacks collected at the indicated time point. Bar, 5 μm.

### Fine structure of LinE in living cells shows that LinEs are axial structures formed on each chromosome

To observe the structure of LinE at higher resolution, we visualized GFP-tagged LinE components in live cells using the super-resolution 3D-SIM system. Fine structure of LinE observed by 3D-SIM in live cells is shown in Figure 3. We found that LinEs in wild-type cells are short, discontinuous filamentous structures along the entire chromosomes in the telomere-clustered horsetail nucleus (Fig 3a, 3b). At karyogamy and the very beginning of the horsetail stage, Rec27-GFP signals were scattered in the whole nuclei. Then, after 30 min in the horsetail stage, a typical LinE structure appeared. At the end of the horstail stage the structure disappeared (Fig. 3c). The LinE number counted from cross section images (Fig. 3d) was about 8 to 12, indicating that LinEs are formed on each chromosome, but do not form in between homologous chromosomes (in this case, if the chromosomes are extended, the maximum number should be six or less at the cross section).

**Fig. 3.**
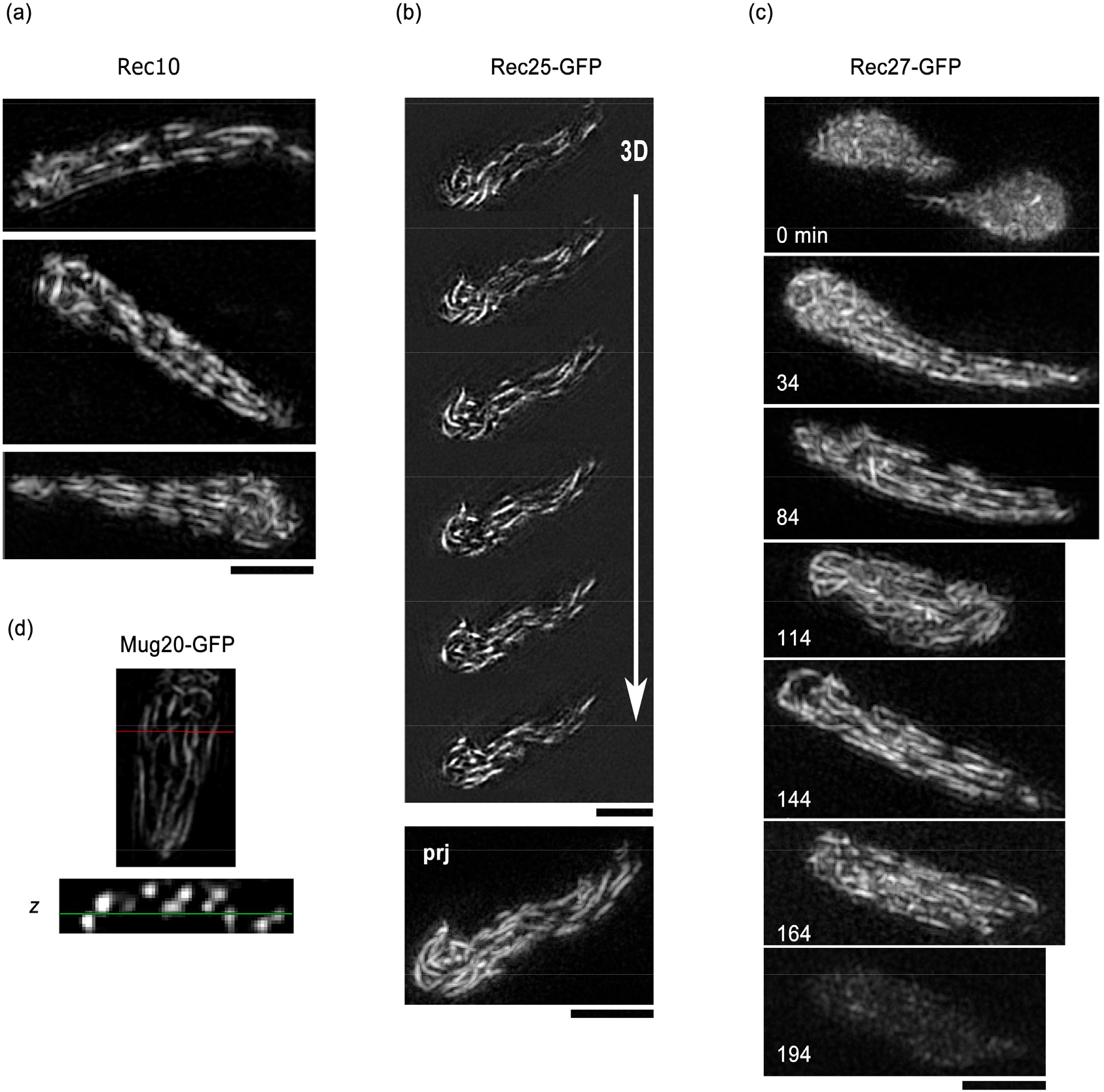
Fine structure of LinE observed in live cells using 3D-SIM. (a) Rec10-GFP in wild-type cells. Three representative projected images from 3D stacks in the horsetail stage are shown. (b) Rec25-GFP in a wild-type cell. Selected continuous Z-focus planes (“3D”) obtained at 0.125 μm focus intervals and their projection (“prj”) are shown. (c) Time lapse observation of Rec27-GFP in a wild-type cell in the entire meiotic prophase. Each image is a projection from 3D stacks collected at the indicated time point. (d) A representative XY plane image and its Z cross section (the XZ plane) image of Mug20-GFP. The red line represents the cross section position and the green line represents the XY plane. Bars, 2 μm.

### Width of LinEs are thinner than cohesin axis

We estimated the apparent width of the LinEs by 3D-SIM imaging analysis (Fig. 4a). The width of the Rec10, Rec25 and Rec27-GFP labeled LinEs were estimated from about 110 nm to 120 nm, which is about the same as the lateral resolution limit of 3D-SIM. The width of LinEs was thinner than the Rec8 width and and histone axis’s width, which were 145 nm and 240 nm, respectively (Fig. 4b, Ding et al. 2015). No statistically significant difference was found between the three LinE components (p >0.025). Cohesin Rec8 axis or the chromosome is significantly wider than LinE (p< 1e-8, Fig. 4b). This suggests that LinEs are formed and restricted on the meiotic cohesin axis.

**Fig. 4.**
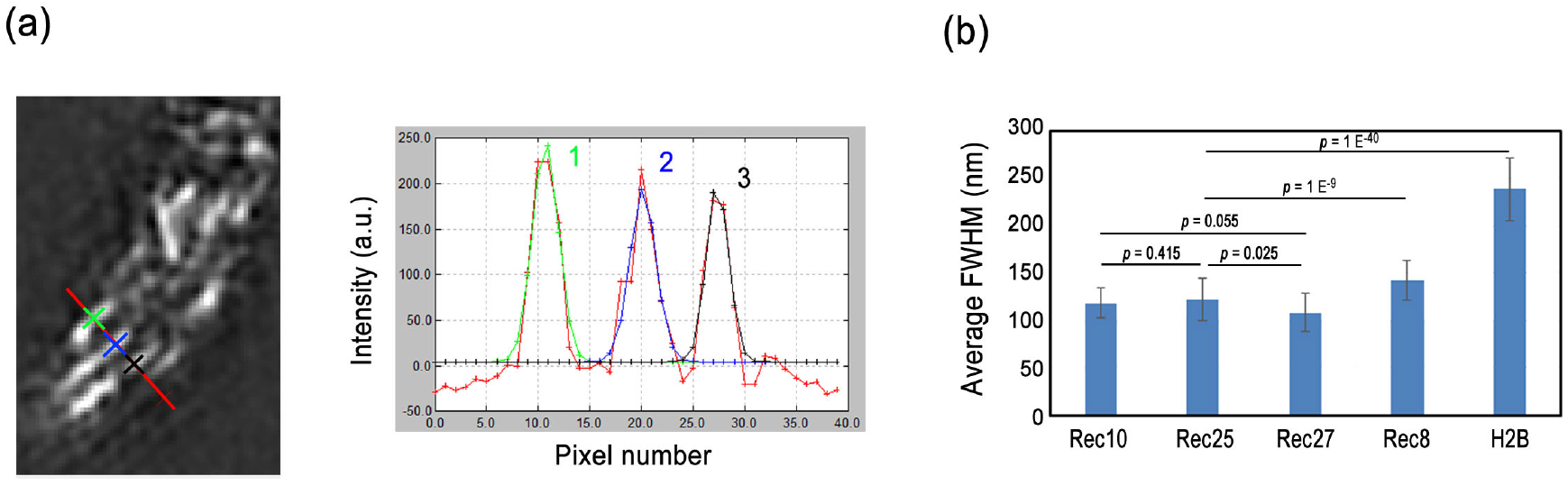
Measurements of the width of LinEs. (a) A line was drawn manually on a section of the image (red line on the left panel), and the pixel intensity along the line was plotted (red line in the right panel). The pixel intensity profile contains three peaks which are numbered. Each separate intensity peak was fitted with a Gaussian profile assuming that the median of the whole 3D stack represents the base intensity with no fluorescence. The full width at half maximum (FWHM) for each numbered peak was calculated from the Gaussian profile. (b) Average FWHM with the standard deviation of Rec10-GFP, Rec25-GFP and Rec27-GFP. The data of Rec8 and H2B are adopted from a previous study (Ding et al., 2015). The *p* values are from two tailed t-test.

**Fig. 5.**
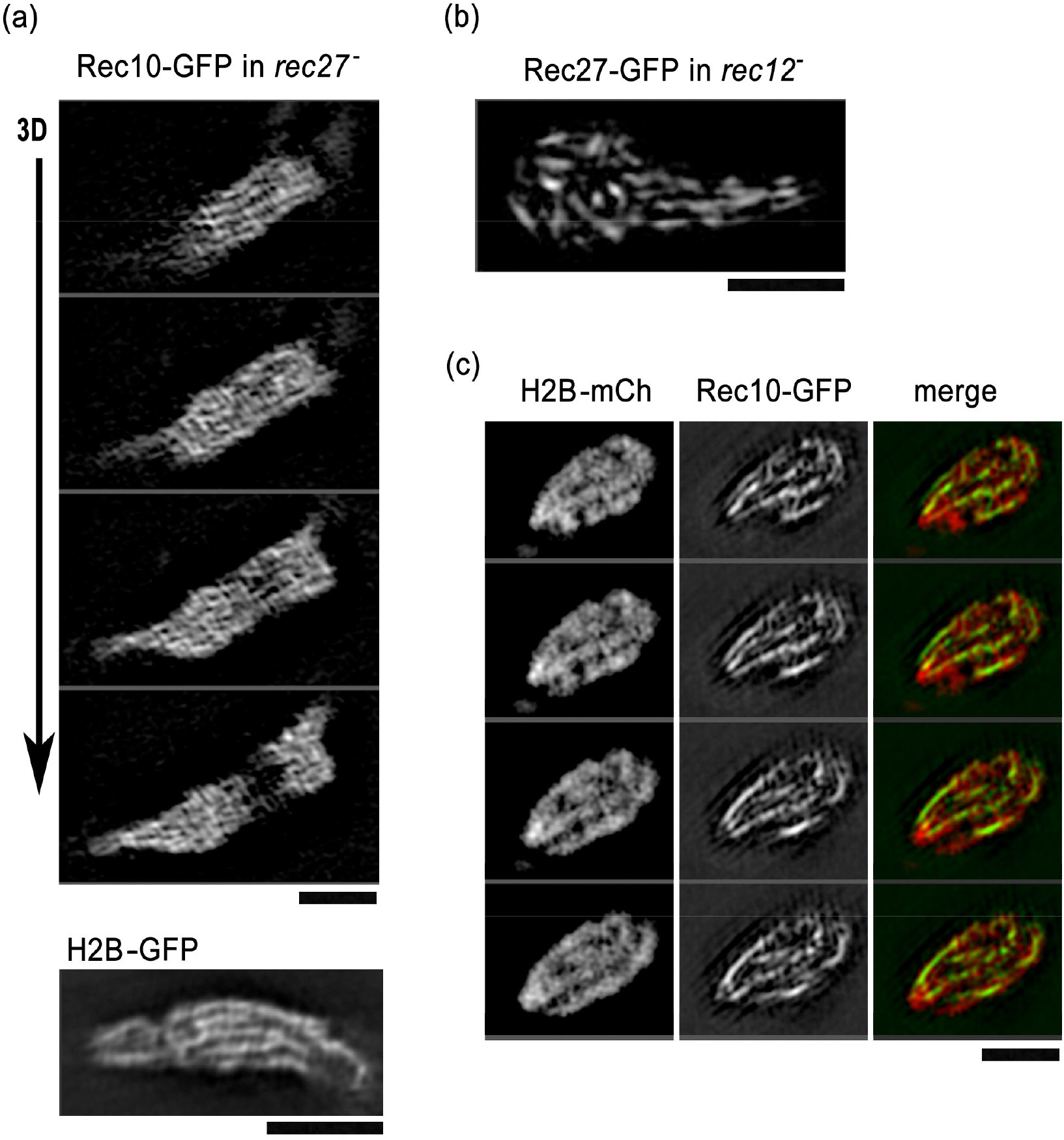
Fine structure of LinE observed in mutant cells using 3D-7SIM. (A) Upper panel: Rec10-GFP in a Rec27 defective cell. Selected continuous Z-focus planes (“3D”) obtained at 0.125 μm focus intervals are shown. Lower panel: A representative single focus plane of H2B-GFP in a wild-type cell. (B) A representative projection image of Rec27-GFP in a Rec12 defective cell. (C) Double labeling of Rec10-GFP and histone H2B-mcherry in a *pds5^-^* cell. Selected continuous Z-focus planes obtained at 0.125 μm focus intervals are shown. Bars, 2 μm.

### 3D-SIM analysis of LinEs in mutant cells

Since in the absence of Rec27, Rec10-GFP showed different labeling pattern compared to that in wild-type cells (Fig. 2a), we further studied its fine structure using 3D-SIM. We found that in Rec27 defective mutant, Rec10-GFP labeled the entire chromosome like in H2B-GFP image (Fig. 5a). This suggests that Rec10 binds to chromatin without other LinEs components. Furthermore, no obvious difference in the shape and dynamics of LinEs was found in a DSB defective mutant rec12-(Fig. 5b) suggesting that DSB formation is unnecessary for LinE formation. In the absence of Pds5, chromosomes are compacted along the longitudinal axis (Fig. 5c, H2B-mCh, Ding et al. 2006; Ding et al. 2015). The LinEs in pds5-mutant show a more continuous appearance (Fig. 5c, Rec10-GFP) compared to that in the wild-type cells indicating the chromosomal axis-localization of LinEs.

### The LinE is a stable structure

It has been found that subunits of SC in *C. elegans* are sensitive to aliphatic alcohols, thus have a liquid-liquid phase separation property (Rog et al. 2017). We have shown that lncRNA–protein complexes in *S. pombe* exhibit phase separation properties, since 1,6-hexanediol treatment reversibly disassembled these complexes and disrupted the pairing of associated loci while the Rec8 axis remained in this treatment (Ding et al. 2019). We then examined that if LinE in *S. pombe* also has liquid droplet properties. We continuously observed LinE proteins in living cells. During live observation, the cells were briefly (5 min) treated with 10% 1,6-hexanediol to resolve the compartments of phase separation. All of the LinE proteins showed little difference by the addition of 10% 1,6-hexanediol for 5 min (Fig.6 a-c). However, a 10 min incubation of the cell in 1,6-hexanediol resulted partial reduction in GFP signal of all the LinE protein-GFP fusion (Fig. 6 d-e). After 10 min incubation in 1,6-hexanediol, the elongated horsetail nuclei shrinked to a small round shape after 5 min incubation in 1,6-hexanediol (the arrows in Fig. 6d-f); the normal horsetail-shaped nucleus did not recover even after 1 hour in culture medium without 1,6-hexanediol, suggesting that the physiology of the cells was disrupted by 1,6-hexanediol treatment longer than 5 min. These results suggested that LinEs are not liquid-liquid phase separated droplets which can be easily resolved in 1,6-hexanediol.

**Fig. 6.**
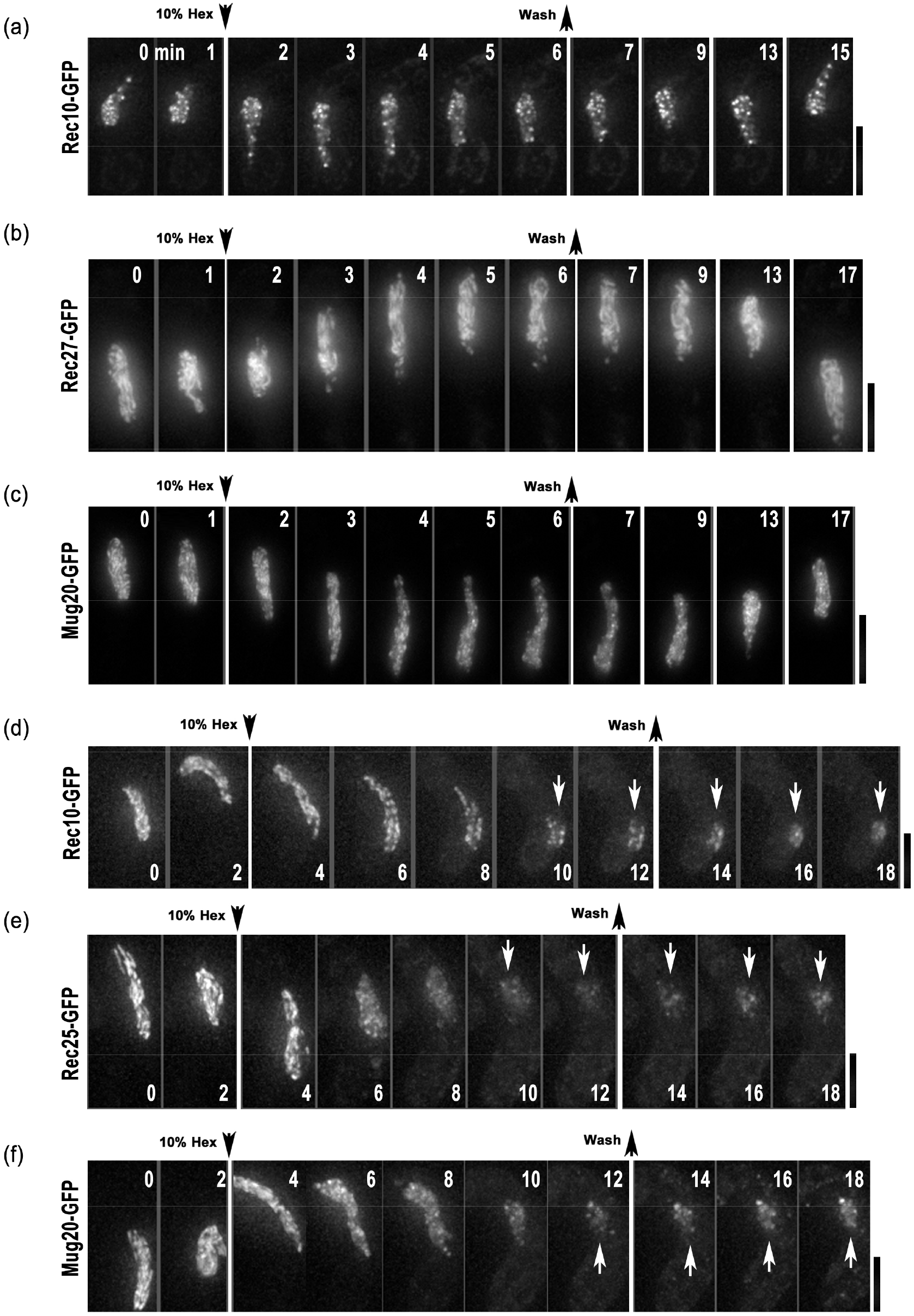
Sensitivity of LinE to 1,6-hexanediol treatment. Time lapse images of LinE proteins in wild-type living cells upon 10% 1,6-hexanediol treatment. 1,6-hexanediol (10%) was added or removed, as indicated by the arrows. (a-c) 5 min treatment. (d-f) 10 min treatment. The numbers indicate the time (minute) of observation. Projected images from 3D deconvolved stacks in the horsetail stage are shown. Arrows indicate the round shaped nucleus. Bars, 5 μm.

Another remarkable feature of the liquid droplet is the ability of rapid fluorescence recovery after photobleaching (FRAP). We then performed FRAP experiments and found that the fluorescence of GFP-tagged LinE proteins did not recover even at 25 sec or more after photobleaching (Fig. 7). Taken together, these data suggest that LinEs are not liquid droplets or liquid crystal but relatively solid and stable proteinous condensates.

**Fig. 7.**
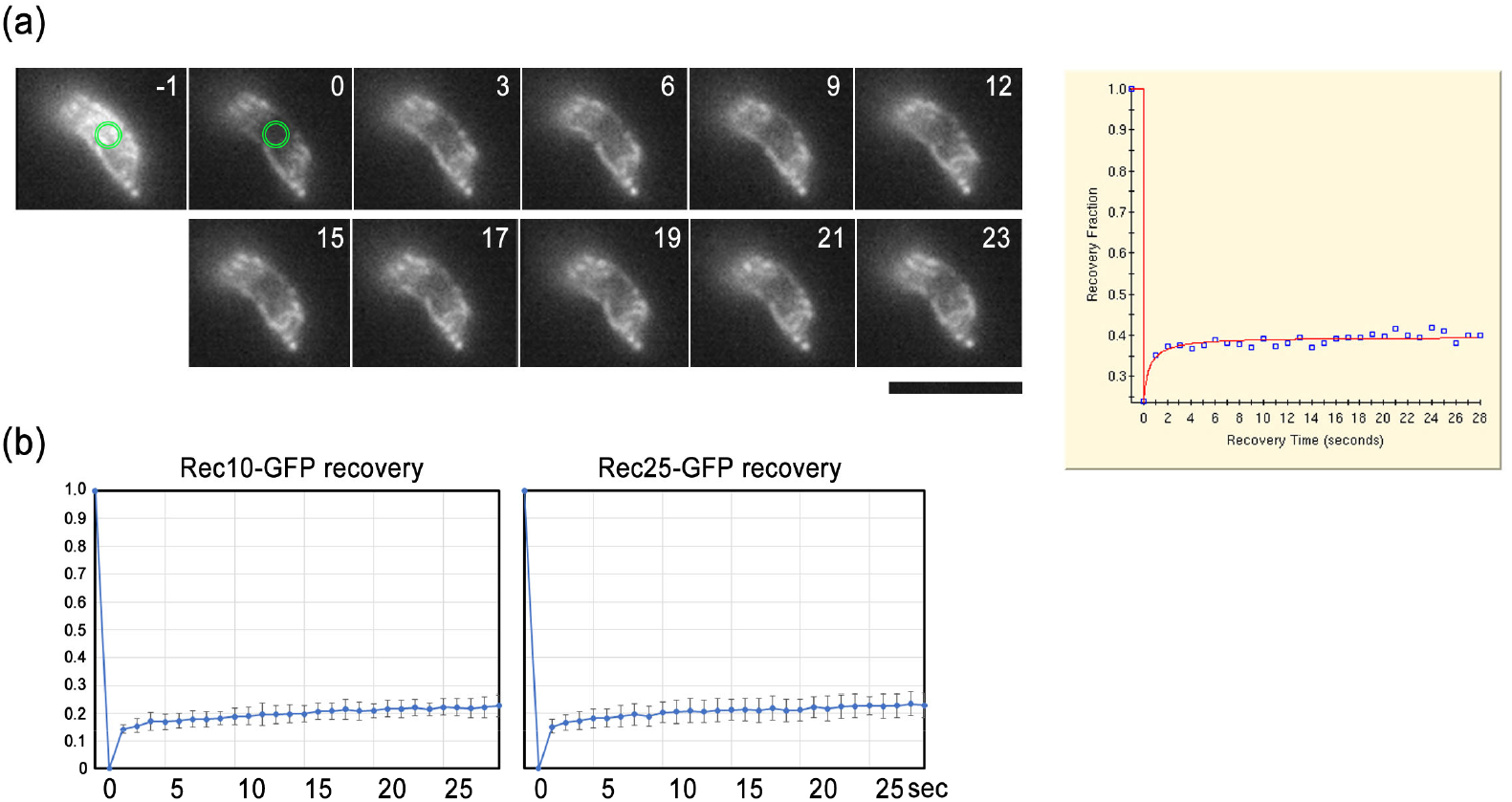
Stability of LinE estimated by FRAP experiment. (a) An example of a FRAP experiment. Rec25-GFP in a wild-type cell before (−1 sec) and after photobleaching (0 sec to 23 sec) at the target spot (green circles). The fluorescence recovery graph (right panel) plots the fluorescence intensity before and after the event. (b) Rec10-GFP and Rec25-GFP recovery after photobleaching. Each graph is an average (+ SD) from 10 cells, Bars, 5 μm.

## Discussion

### Hierarchy of LinE establishment

In this study, we followed the dynamics and interdependency of LinE proteins in living cells during the entire meiosis. Although Rec10, Rec25, Rec27 and Mug20 are all the essential components of LinEs, Rec10 could bind to the chromosomes without the rest of LinE components examined in this study (Fig. 2a-c, Fig. 5a), while other LinE components could not bind to the chromosomes in the absence of Rec10 (Fig. 2d) (Fowler et al. 2013). It has been found that the C-terminus of Rec10 binds to CK1 phosphorylated meiotic cohesin Rec11 (Sakuno and Watanabe 2015). Thus, Rec10 may plays a major role in linking LinEs with meiotic cohesin and the meiotic chromosome. Furthermore, using 3D-SIM, we found that LinEs are uniform in width and even thinner than the width of the meiotic cohesin axis (Fig. 3, Fig. 4b). This implies that the formation of LinEs is more constrained to the meiotic chromosome axis then the cohesin complex, or implies a lesser abundancy of LinE proteins compared to the cohesin proteins.

The typical short filamentous structure forms only when all four essential LinE components are present. All LinE components show similar dynamics that they appear in the nucleus from karyogamy and disappear at the end of the horsetail stage, before meiosis I (Figure 1a, 1b). Without the formtion of LinEs, Rec10 and Mug20 were regulated differently. They persistently stay in the nucleus even after meiosis I (Figure 2a, 2b), which is similar to the dynamics of meiotic cohesins (Ding et al. 2006; Sakuno and Watanabe 2015). These data suggest that LinE proteins are regulated differently upon forming the LinE filamentous complex. Since LinEs are critical for the formation of meiotic recombination hotspots (Fowler et al. 2013), the LinE components are probably regulated together with other protein factors that function as recombination machinery.

### Structure and dynamics of LinEs in living cells revealed by 3D-SIM imaging

Using 3D-SIM, we obtained images of LinEs in living cells with super high resolution at 120 nm in lateral and 350 nm in the axial direction. Our results confirmed that LinEs in *S. pombe* represent an axial structure that colocalizes with the chromosomal axis (Bahler et al. 1993; Lorenz et al. 2004; Lorenz et al. 2006), but not the central elements, which link the axial or lateral elements into SCs. We have not observed so called networks or thick bundles or dots when tracked the entire meiotic prophase in the live cells, which suggests that those images from EM analyses can be artifacts created in the sample preparation (Lorenz et al. 2004; Molnar et al. 2003).

### Stability of LinEs

It was shown that Rec27 and *C. elegans* SC protein SYP-2 share DNA sequence similarity at their coiled-coil domain (Fowler et al. 2013) suggesting that LinEs may have similar properties with the central element component of SC. The central element proteins of SC, except the chromosome axis, in *C. elegans, S. cerevisiae* and *Drosophila melanogaster* can be dissolved in the presence of 1,6-hexanediol suggesting that the central elements of SC form through liquid-liquid phase separation (Rog et al. 2017). We tested if LinEs have a liquid-liquid phase separation property by treating it with 1,6-hexanediol or through FRAP experiment. However, we found a very solid and stable property of LinEs (Fig. 6a-c). This stable property is consistent if LinEs represent axial elements along the chromosome axis, but do not represent transverse elements in between homologous chromosomes. No SC central element-like transverse element has ever been found in *S. pombe*. The only structures observed in between homologous chromosomes are lncRNA/RNA-transcription termination protein complexes, which are liquid-liquid phase separation droplets that play an essential role in the recognition and pairing of homologous chromosomes (Ding et al. 2019). Since the liquid crystalline-like, transverse elements of SC may play a direct role in crossover interference (Rog et al. 2017), and no crossover interference exists in *S. pombe*, LinEs on the chromosome axis may play roles in crossover control in a different way from that of SC (Loidl 2006; Yamada et al. 2017). Further research on SC-less organisms will provide more insights into the recombination control specifics for meiosis.

## Materials and Methods

### Strains and culture

The *S. pombe* strains used in this study are listed in Table 1. *S. pombe* standard culture media YES, ME, and EMM2-N were used for routine culture, meiosis induction, and live observation, respectively. Strains for live cell observation were constructed as follows: GFP or mCherry tagging and gene deletions were created using a PCR-based gene-targeting method (Bahler et al. 1998). In GFP or mCherry tagging, the open reading frame of GFP or mCherry was integrated at the C-terminal end of the endogenous gene locus in the genome. Microfluidic yeast plates (CellASIC) were used for cell culture and 1,6-hexanediol treatment on the microscope stage.

**Table 1.** Strain list

| Strain | Genotype |
| --- | --- |
| YY365 | *h<sup>90</sup> ade6-216 leu1-32 lys1-131 ura4-D18*<br>*rec10<sup>+</sup>::GFP-kan<sup>r</sup>* |
| YW046 | *h<sup>90</sup> ade6-216 leu1-32 lys1-131 ura4-D18*<br>*rec25<sup>+</sup>::GFP-kan<sup>r</sup>* |
| YW047 | *h<sup>90</sup> ade6-216 leu1-32 lys1-131 ura4-D18*<br>*rec27<sup>+</sup>::GFP-kan<sup>r</sup>* |
| YY965 | *h<sup>90</sup> ade6-216 leu1-32 lys1-131 ura4-D18*<br>*mug20<sup>+</sup>::GFP-kan<sup>r</sup>* |
| YW045 | *h<sup>90</sup> ade6-216 leu1-32 lys1-131 ura4-D18*<br>*rec10<sup>+</sup>::GFP-kan<sup>r</sup> Δrec27::hyg<sup>r</sup>* |
| YY974-6A | *h<sup>90</sup> ade6-216 leu1-32 lys1-131 ura4-D18*<br>*mug20<sup>+</sup>::GFP-kan<sup>r</sup> Δrec10::ura4<sup>+</sup>* |
| YW048 | *h<sup>90</sup> ade6-216 leu1-32 lys1-131 ura4-D18*<br>*rec25<sup>+</sup>::GFP-kan<sup>r</sup> Δmug20::hyg<sup>r</sup>* |
| YY044 | *h<sup>90</sup> ade6-216 leu1-32 lys1-131 ura4-D18*<br>*rec10<sup>+</sup>::GFP-kan<sup>r</sup> Δrec25::hyg<sup>r</sup>* |
| YY940 | *h<sup>90</sup> ade6-216 leu1-32 lys1-131 ura4-D18*<br>*rec10<sup>+</sup>::GFP-kan<sup>r</sup> Δmug20::hyg<sup>r</sup>* |
| YY362 | *h<sup>90</sup> ade6-216 leu1-32 lys1-131 ura4-D18*<br>*rec10<sup>+</sup>::GFP-kan<sup>r</sup> rec12-152::LEU2* |
| YW275 | *h<sup>90</sup> ade6-216 leu1-32 lys1-131 ura4-D18 Δpds5::LEU2*<br>*rec10<sup>+</sup>::GFP-kan<sup>r</sup> aur1<sup>r</sup>::[hta1<sup>+</sup>-htb1<sup>+</sup>-mCherry]* |

### Microscopies

For deconvolution microscopy, a DeltaVision Elite microscope (GE Healthcare, Buckinghamshire, UK) with an objective lens 60X PlanApo NA 1.4 Oil SC (Olympus) set up in a temperature-controlled room at 26°C was used. A set of images from 15 focal planes with 0.3 μm intervals was taken every 5 min. Data analysis was carried out by using SoftWoRx software on the DeltaVision system.

For 3D-SIM imaging, we used a DeltaVision|OMX microscope version 3 (GE Healthcare) with an objective lens 100X UPlanSApo NA1.40 Oil (Olympus, Tokyo, Japan). For live cell SIM, cells in EMM2-N attached to glass-bottom dishes coated with lectin were imaged with immersion oil with a refractive index of 1.522. Live cell SIM reconstruction was performed by using the softWoRx software (GE Healthcare) with a wiener filter constant of 0.012. To cover the entire nucleus, a set of 17 optical sections were taken at 0.125 μm focus intervals. For simultaneous observations of H3-mCherry and Rec10-GFP, a set of 9 optical sections were taken. Our custom software was used for the correction of chromatic aberrations, and camera alignment (Matsuda et al. 2018). The LinE width was measured by 1D Gaussian fitting and FWHM was obtained by 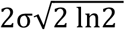, where σ is the standard deviation of the Gaussian distribution. We selected cells that were undergoing horsetail movements irrespective of the prophase stage and had fluorescent signals strong enough to perform the 3D-SIM analysis.

Photobleaching and recovery experiments were conducted using the Photokinetics function of DeltaVision|OMX with the same lens described above. We used 488 nm laser, 10% T, and duration of 0.01 sec to bleach the GFP fluorescence signal. The FRAP data analysis was performed by using the PK analysis function of the softWoRx software with the two components model.

## Acknowledgments

This research was funded by KAKENHI grants from MEXT of Japan (19K06503 to DQD., 19H03202 to AM., JP17H01444 and JP18H05533 to YH).

